# Specializations in Tail Anatomy of the Lesser Egyptian Jerboa (*Jaculus jaculus*) Compared with the Mouse and Rat

**DOI:** 10.64898/2026.06.21.733634

**Authors:** Juri A. Miyamae, Talia Y. Moore

## Abstract

Mammal tails have long been recognized for their diversity of morphological form and function, however, there remains a substantial gap between the motivation to understand and emulate the various performance functions of the tail and what is known about tail anatomy. In this study, we were motivated to discover the anatomical foundations of the fast, whipping motions of the tail of the lesser Egyptian jerboa (*Jaculus jaculus*), which may aid in the quick changes of direction as the animal escapes from predators using ricochetal bipedal hopping. We employed microCT scans, dissections, and museum data to describe the musculoskeletal anatomy of the jerboa in comparison with the laboratory mouse (*Mus musculus*) and rat (*Rattus norvegicus*). While many aspects of tail anatomy are conserved across these species, the jerboa does possess unique characteristics such as an extremely long tail arising from caudal vertebral elongation, development of extensive dorsal musculature differentiated into lateral and medial components to increase points of skeletal attachment, and a novel anatomical feature – the bi-lobed cranial transverse process – which serves as a supernumerary dorsal tendon attachment site and possible brace to protect the ventral tendons and intrinsic muscles for a section of caudal vertebrae which likely experiences high mechanical stress.

## INTRODUCTION

Humans are relatively unusual in lacking external tails, the ubiquitous “fifth limb” found across all tetrapods. Just amongst mammals, tails have diversified into a remarkable range of morphology and functions which have been investigated from multiple disciplinary perspectives. Morphological descriptions of the tail can be as subtle as assessments of relative lengths and tufting patterns to distinguish between closely-related species (e.g., Heritage et al. 2020), or capture significant evolutionary transformations such as tail-loss in Hominoidea in correlation to the emergence of human bipedalism (Machnicki et al. 2016; Williams and Russo 2015) and the development of fluked tails powering the swimming of cetaceans as they transitioned into fully aquatic life (e.g., Buchholtz 2001, 2007; Buchholtz and Gee 2017; Gillet et al. 2024).Functionally, tails encompass roles as varied as intraspecific and interspecific communication (e.g., Blank 2018; Cafazzo and Natoli 2009; Caro et al. 1995; De Boer et al. 2013; Haulenbeek and Katz 2011; Hennessy et al. 1981; Hersek and Owings 1993; Palagi 2009; Walker-Bolton and Parga 2017), thermoregulation in response to both heat and cold (e.g., Bennett et al. 1984; Dawson and Keber 1979; Muchlinski and Shump 1979; Škop et al. 2020; Stryjek et al. 2021), prey capture in bats (e.g., Webster and Griffin 1962), fat storage (e.g., Aleksiuk 1970; Blanco et al. 2022; Booth and Connolly 2008; Farhadi et al. 2021; Gordon and Hall 1995; Harris 1987; Lemelin and Schmitt 2004), carrying of nesting materials among marsupials and platypus (e.g., Dalloz et al. 2012; McManus 1970; Macrini 2004; Soares et al. 2024; Stodart 1966; Thomas et al. 2018), aiding parental care through protecting and transporting clinging young (e.g., Bertassoni et al. 2025; Griffiths 1988; Wimsatt 1960; Zhang et al. 2015), as bludgeoning weapons (e.g., Alexander et al. 1999; Arbour and Zanno 2020), giving potential predators the slip through tail skin sloughing or autotomy in small rodents (Dubost and Gasc 1987; Juškaitis 2006; Layne 1972; Seifert et al. 2012; Shargal et al. 1999), tactile sensation (e.g., Hajyahia et al. 2025; Hickman and Brown 1973; Organ et al. 2011), warding off pests and parasites (e.g., Barros et al. 2024; Matherne et al. 2018; Mooring et al. 2007), and even expressing group-specific learned behaviors such as “tail walking” in bottlenose dolphins (e.g., Bossley et al. 2018). However, most research interest has focused on the locomotory role of the mammalian tail as either a dynamic counterbalance, especially while traversing unstable substrates (e.g., Alexander and Vernon 1975; Buck et al. 1925; Wada et al. 1993; Walker et al. 1998; Young et al. 2015, 2021), or an inertial appendage to reorient the body (e.g., Fukushima et al. 2021; Patel et al. 2016; Schwaner et al. 2021). Behavioral observations, experimental data, and biomechanical modeling of tails across vertebrates – not just mammals – have found their place in bioinspired engineering, which has sought to emulate the counterbalancing and inertial maneuvering functions for both robots and human wearers (e.g., Anwar et al. 2024; Briggs et al. 2012; Huang et al. 2024; Kohut et al. 2013; Libby et al. 2012; Nabeshima et al. 2019; Storms and Tilbury 2016).

The basic organization of the tail is highly conserved. All vertebrates – even “tailless” ones – possess an external tail at some point in their development. This post-anal extension of the axial column is a feature not only of vertebrates, but a synapomorphy at a deeper ancestral node encompassing all chordates. Like all other vertebrates, the mammalian tail is supported by caudal vertebrae and mobilized by bilateral epaxial (i.e., dorsal) and hypaxial (i.e., ventral) musculature (Liem et al. 2001). Mammals have modified this basic architecture through the evolution of morphologically distinct caudal vertebrae – presumably as part of a pattern of increasing vertebral regionalization and specialization that began among non-mammalian synapsids (Jones et al. 2018; Jones et al. 2019) – as well as greater decoupling of the caudal from the hindlimb musculature (Haines 1935; Howell 1938) and shifting from a muscle-dominated to a more lightweight, tendon-dominated tail (Howell 1938; pers. obs. JAM/TYM).

Though the mammalian tail presents such distinctive features, anatomical representations have been scarce and dispersed across the literature. Most anatomical studies detailing musculoskeletal morphology are dedicated to single species descriptions, e.g., western grey kangaroo (*Macropus fuliginosus;* Dawson et al. 2014), grizzled tree kangaroo (*Dendrolagus inustus;* Vrolik 1857), aardvark (*Orycteropus afer;* Endo et al. 2013), domestic dog (*Canis familiaris;* Miller et al. 1964), domestic cat (*Felis catus;* Crouch 1969), beaver (*Castor canadensis;* Mahoney and Rosenberg 1981), rat (*Rattus norvegicus*; Brink and Pfaff 1980; Hori et al. 2011), and mouse (*Mus musculus;* Shinohara 1999). In contrast, comparative anatomies suited for phylogenetic or functional interpretation are relatively few, such as a study on the general anatomy allowing enrolling among Indian hedgehogs that includes tail musculature (*Paraechinus micropus micropus* and *Hemiechinus auritus collaris;* Gupta 1965) and an investigation into the musculoskeletal morphologies associated with tail prehensility in various New World primates (German 1982; Lemelin 1995; Organ 2010). Veterinary anatomical textbooks (Ashdown et al. 2011; Aspinall and Cappello 2015; Cochran 2011; Done et al. 2009; Hermanson et al. 2020; König and Liebich 2020; Singh 2018) and vertebrate dissection guides (Evans and De Lahunta 2017; Homberger and Walker 2004; Kardong and Zalisko 2012; King and Custance 1982) tend to present only limited written descriptions (if at all) of tail anatomy and illustrative diagrams that frequently fade out at the tail, offering literally truncated information about the structure and organization of this appendage. Therefore, despite recognition of the varied and vital importance of tails in the daily lives of mammals, as well as their rich potential for practical applications in engineering, there is a disproportionate paucity of studies exploring the musculoskeletal anatomical foundations that generate the functional diversity of mammalian tails. Without robust anatomical knowledge, necessarily set within a comparative framework, our ability to identify the core mechanical principles driving form-function relationships will be inherently limited.

To begin filling this gap, we present a comparative anatomical case study of the tail of the lesser Egyptian jerboa (*Jaculus jaculus*; Rodentia, Dipodidae) and two common laboratory rodents, the mouse (*Mus musculus*; Rodentia, Muridae) and rat (*Rattus norvegicus domestica*; Rodentia, Muridae) (Figure 1). We use data collected primarily from dissections and microCT scans to produce descriptions of gross morphology, osteology, myology, and maps of tail tendon insertions. These findings are used to hypothesize key anatomical features associated with different tail functions and demonstrate aspects of intraspecies variation.

**Figure 1.**
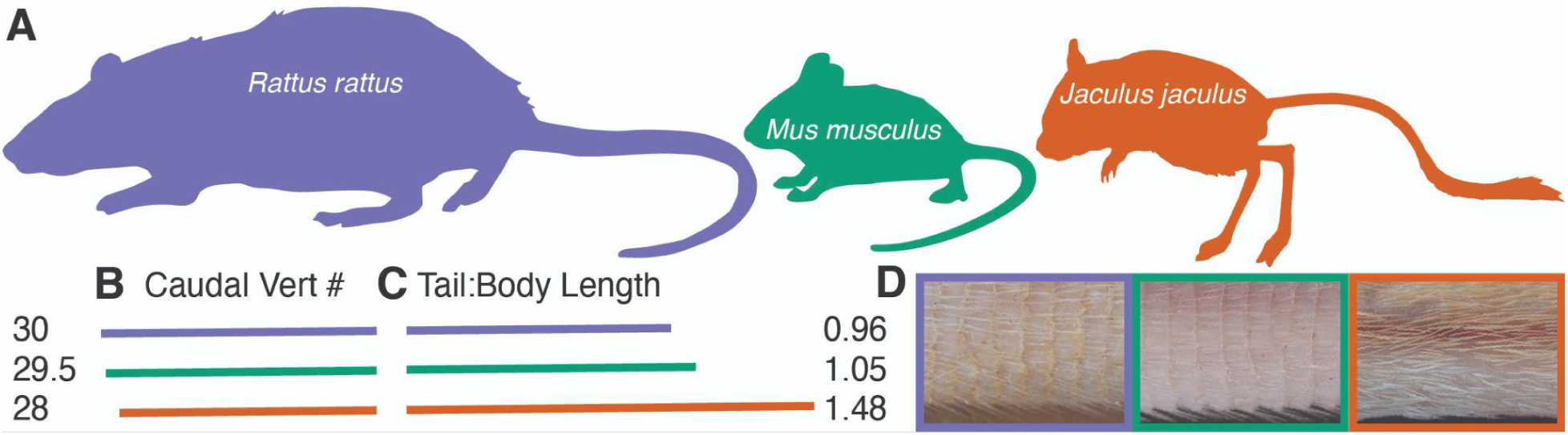
Gross morphology of the three rodents examined. A) Body size differs greatly, but B) caudal vertebra number does not vary much among the three rodents. C) Tail to body length ratio is significantly higher in jerboas than in rats and mice. D) Photographs show detail of the skin and hair on each tail. Silhouettes are public domain images by Ferran Sayol (rat), Jiro Wada (mouse), and Margot Michaud (jerboa), obtained via PhyloPic.

Our selection of rodent species for comparison aims to identify which anatomical characters of tails may have evolved in tandem with locomotory specializations. The jerboa is a bipedal hopper that escapes from predators by employing high-speed ricochetal flight paths over shifting desert sands. This entails erratic and rapid changes in direction (Moore et al. 2017), which may be aided by the whipping movements of its long tail. A similar mechanism has been demonstrated in kangaroo rats (*Dipodomys spp*.; Rodentia, Heteromyidae), a convergently bipedal desert rodent that uses wide swings of its tail to reorient its body mid-air in the yaw-plane when leaping to escape rattlesnake strikes (Chu et al. 2024; Schwaner et al. 2021). In contrast, laboratory mice and rats are frequently regarded as generalist quadrupeds or locomotor generalists, especially in establishing comparative experimental and clinical models of locomotion (Charles et al. 2016; Polly 2007; Schmidt and Fischer 2011; Sears et al. 2006; Wright et al. 2022). Our investigations show that while many anatomical features remain conserved between jerboa, mice, and rats, the locomotory specializations of the jerboa appear to be most correlated with gross tail proportions, caudal vertebrae proportions and morphology, muscle differentiation, and the relative distribution of dorsal versus ventral tail muscle masses.

## MATERIALS & METHODS

Osteological descriptions were taken from microCT scans obtained from MorphoSource (morphosource.org) and processed using Materialise Mimics Version 24.0.0.427 (Materialise NV, Leuven, Belgium; materialise.com). Scans were selected for completeness of the caudal vertebrae series, typically found in scans of whole body soft tissue specimens, where caudal elements are less likely to be lost and/or disarticulated. Specimen information, MorphoSource media ID number, microCT scanning facility, and microCT scanning parameters are as follows:

### Mus musculus

TMM.M.8671; whole body scan; media ID 000606487; Varian Medical Systems (Bio-Imaging Research, Inc.) ACTIS CT scanner with FeinFocus x-ray source at University of Texas High-Resolution X-ray Computed Tomography Facility; pixel spacing x, y = 0.0371 and z = 0.0842 mm; volt n/a, amp n/a.

### Rattus norvegicus

UMMZ:mammals:167022. MorphoSource media ID 000082216. Scan data acquired with Nikon Metrology XT H225 ST at University of Michigan Museum of Zoology, Research Museums Center. Scanned at n/a kV and n/a μA. Scan resolution 0.08886 mm.

### Jaculus jaculus

UMMZ:mammals:120204. MorphoSource media ID 000064399. Scan data acquired with Nikon Metrology XT H225 ST at University of Michigan Museum of Zoology, Research Museums Center. Scanned at 85 kV and 200 μA. Scan resolution 0.07624 mm.

For better comparative visualization of the tail vertebrae, each bone was segmented as a separate region of interest (ROI) in Materialise Mimics, exported as a STL file, and then imported into Blender (version 4.4.3, Blender Foundation, Amsterdam, Netherlands) to align all the vertebrae along a common cranial-caudal axis and measure vertebral dimensions. Vertebral centrum length (*L_c_*) was measured along the central cranial-caudal axis of each vertebrae. *L_c_* was used to plot centrum length versus vertebral position. Maximum vertebral length, width, and height (*L_max_, W_max_, H_max_*) encompass the entire bone, including processes that may extend cranially past the proximal edge of the centrum. We constructed a bounding box for each aligned vertebra and ensured it was aligned with the global axes. Because bounding box dimensions are defined by the maximum dimensions of an object in each axis, the bounding box length, width, and height correspond to *L_max_, W_max_, H_max_*, respectively.

Gross external tail morphology and soft tissue anatomy of the muscles and tendons were examined through microdissections of both 10% buffered formalin fixed and unfixed rodent carcasses: mouse n = 4 consisting of 1 female, 1 male, 2 unknown sex, with 1 fixed and 3 fresh specimens; rat n = 1 fresh specimen of unknown sex; jerboa n = 4, consisting of 2 female, 1 male, 1 unknown sex, with 2 fixed and 2 fresh specimens. All specimens are laboratory-raised animals under IACUC-approved protocols from AAALAC-accredited facilities (University of Michigan Ann Arbor, University of California San Diego, and Harvard University). While we endeavored to represent “standard” adult tail anatomy, information on age, sex, and strain were not always available for the mouse and rat specimens in this study, as these were salvaged carcasses from a training core. However, these were all healthy animals with no outstanding gross musculoskeletal anomalies and we have mitigated potential sampling issues through the dissection of multiple individuals for each species. All jerboas were laboratory-bred wildtype. Dissections were performed using a Leica M60 stereomicroscope equipped with an IC90E Integrated CMOS Microscope Camera. Additional images were taken with a Nikon D500 with a 50 mm f/1.8 lens augmented with Kenko extension tubes and lighting provided by a Godox Ring72 Macro LED ringlight.

Lengths were measured using either a Helios dial caliper (d = 0.05mm) or office ruler (d = 1 mm). Body lengths were measured from the tip of the snout to the base of the tail. Tail lengths were measured at the base of where the tail bends at the sacrocaudal joint to the fleshy tip of the tail. These lengths were used to calculate tail:body length ratios. Muscle lengths were measured while wet and along their longest linear dimension. Tendon lengths were likewise measured while wet, gently straightened out on a flat surface while avoiding excessive stretching, and measured from the tip of the aponeurosis – either while still attached to the muscle or carefully scraped off the muscle – to the free end where the tendon was cut close to the insertion point.

Weights were measured using an Accuteck Digital Postal Scale (d = 1 g / 0.1 oz), Smart Weigh GEM20 digital jewelry scale (d = 0.001 g), or OHAUS Navigator NV323 portable balance (d = 0.001 g). Total body weights were collected in our lab when feasible from intact specimens or taken from necropsy reports or data provided upon receipt of the specimen. Muscle weights were taken from excised muscles cleaned of tendons, fat, and connective tissue. In some samples, both wet and dry weights were taken. Wet weights were taken immediately after collection and dabbed with a paper towel to remove excess moisture before weighing. DDry weights were taken from muscle that have been air dried at room temperature for a minimum of 72 hours. Weights were collected in triplicate and values averaged.

Tail:body length ratios from our sample of captive rodents were compared with measurements from wild populations collected by natural history museums and made accessible online via Global Biodiversity Information Facility (www.gbif.org). Search for species name *Occurrences* was further refined with the search terms: *Search all fields* = “tail” and *Basis of record* = “Preserved specimen.” Measures are typically reported in the *Dynamic properties* or *Measurement or Fact* field for each entry. If only “total length” was reported, then “tail length” was subtracted from that value to calculate body length. We collected measurements from n = 10 individuals from each species, with an equal number of males and females, if available. To facilitate a more diverse sampling that does not overrepresent a particular collection locality or institution, measurement data were collected at a predetermined interval depending on the number of entries available (e.g., for *Mus musculus* and *Rattus norvegicus*, first entry meeting data completeness and sex sampling criteria on every 10^th^ page of search results).

## RESULTS

### Gross morphology

The mouse, rat, and jerboa all possess long, gently tapering, and slender tails (*Figure 1A*). In superficial appearance, they all have stereotypically “naked” rodent tails devoid of the pelage hairs that cover the body. However, the underlying skin of mouse and rat tails is visible beneath a scant covering of short, bristle-like hairs which have been characterized as a specialized hair type with its own unique morphogenesis (*Figure 1D*, Alibardi 2004; Duverger and Morasso 2009).

Mouse tail:body length ratio from our sample is 0.96 ± s.d. 0.09 (n = 4, *Figure 1C*, green); the tail:body length ratio of wild *Mus musculus* collected from data reported from museum specimens in GBIF is 0.96 ± s.d. 0.11 (n = 10). The tail is approximately circular in cross-section. The skin of the tail is visible and shows parallel, transverse annulations regularly spaced approximately 0.9 mm apart (*Figure 1D*, green). The number of these tail rings averages 180, ranges from 142 – 220, but shows low correlation between the number of annulations and tail length (Fortuyn 1928). Short bristles of hair cover the tail and grow in clumps of three or four, a grouping pattern also observed in previous studies (Chang et al. 2014; Duverger and Morasso 2009). Hairs emerge from beneath the epidermal folds in the interscale regions (Duverger and Morasso 2009) that form the distal edge of the annulations. There are no marked morphological differences between the dorsal and ventral surface, or along the length of the tail.

Rat tail:body length ratio from our sample is 1.05 (n = 1; *Figure 1C*, purple); the tail:body length ratio of wild *Rattus norvegicus* from data reported in GBIF is 0.84 ± s.d. 0.08 (n = 10). The tail is especially heavy and robust compared to the mouse and jerboa. The cross-section is approximately circular. Hair coverage is the most sparse of the three species examined, revealing raised and easily visible annulations, approximately 1.1 – 1.3 mm apart. There are an estimated 210 (Ugwu 1987) to 250 annulations (Bao 1995). In addition, the annulations are further segmented to form distinct scales with a ragged free distal edge that overlaps the subsequent ring of scales (*Figure 1D*, purple). While we observed a relatively uniform distribution of hair bristles emerging from beneath the scales, other authors have noted groups of three hairs per scale (Bao 1995) or a range of one to seven hairs per scale depending on location, with greater numbers of hair more prevalent proximally (Erickson 1931). The rat tails we observed had a patchy covering of a yellow waxy secretion, which has also been noted in previous descriptions, but without further comment on the composition of this material or any potential functional hypotheses (Bao 1995; Ugwu 1987). There is relatively little morphological difference in regions of the tail, except that the ventral tail hairs appear slightly longer than the dorsal.

Jerboa tail:body length ratio from our sample is 1.69 ± 0.02 (n = 3; *Figure 1C*, orange); the tail:body length ratio of wild *Jaculus jaculus* from data reported in GBIF is 1.66 ± s.d. 0.08 (n = 8). These proportions far exceed that of the mouse and rat, where the tail is approximately the same length or slightly shorter than the body. The tail is approximately square in cross-section, but more circular in particularly well-fed individuals with accumulated subcutaneous fat. Unlike the mouse and rat, the tail is covered with relatively long hairs that largely obscure the skin beneath (*Figure 1D*, orange); however, removal of the hair reveals only very faint and irregular annulations, approximately 0.5 – 1.0 mm apart. Despite this diminished epidermal sculpturing, the regular placement of hair follicles follows a vaguely ring-like pattern similar to that seen in mice and rats. Unlike the laboratory strains of mice and rats we examined that have a more or less uniform coat color, jerboa pelage consists of a fawn or sandy brown dorsum and white ventrum – a color scheme that continues with the tail hairs. The division between the brown dorsal hairs and white ventral hairs is approximately demarcated by the lateral tail veins that are prominently visible beneath what is apparently a relatively thin and delicate skin. Finally, the jerboa tail is distinguished by a terminal tuft of black and then white hair. The black portion often does not completely encircle the tail, covering only the dorsal and lateral aspects. The white portion consists of long hairs that extend beyond the fleshy tip of the tail.

Please refer to Supplemental Materials, *Table 1* for measurements collected for this study and *Table 2* for measurements and specimen data gathered from GBIF.

### Osteology of the Caudal Vertebrae

Despite the remarkable tail length in jerboas, all three examined rodent species possess similar numbers of caudal vertebrae: mouse mean = 30.4 (range = 29 – 32; n = 5), rat mean = 29.5 (range = 29 – 30; n = 2), and jerboa mean = 27.8 (range = 27 – 28; n = 5)(*Figure 1B*; refer to Supplemental Materials, *Table 3* for data). Therefore, the length of the jerboa tail is generated through the elongation of each individual caudal vertebra, instead of increasing the number of vertebrae. *Figure 2H* shows centrum length versus anterior to posterior vertebral position, displaying an overall “crescendo-decrescendo” (Shinohara 1999a) pattern in all three species, where each vertebral centrum is longer than its antecedent, rising to a peak maximum length, and then decreasing in length with each successive vertebrae to the end of the tail. Both the mouse and rat show similar shapes in this crescendo-decrescendo curve, though the larger rat does demonstrate greater absolute centra lengths. Maximum centrum length in the mouse is found in the 7^th^ vertebral position, i.e., caudal vertebra (Cd) 7, with a length of 4.08 mm. Maximum central length in the rat is found in Cd 8 with a length of 7.77 mm. The longest vertebral centrum in the jerboa occurs in a similar position at Cd7, but with a length of 9.46 mm. In contrast to the mouse and rat, the jerboa crescendo in centra lengths rises rapidly to a sharp peak, superseding the lengths in the rat from vertebral position 6 to 18.

**Figure 2:**
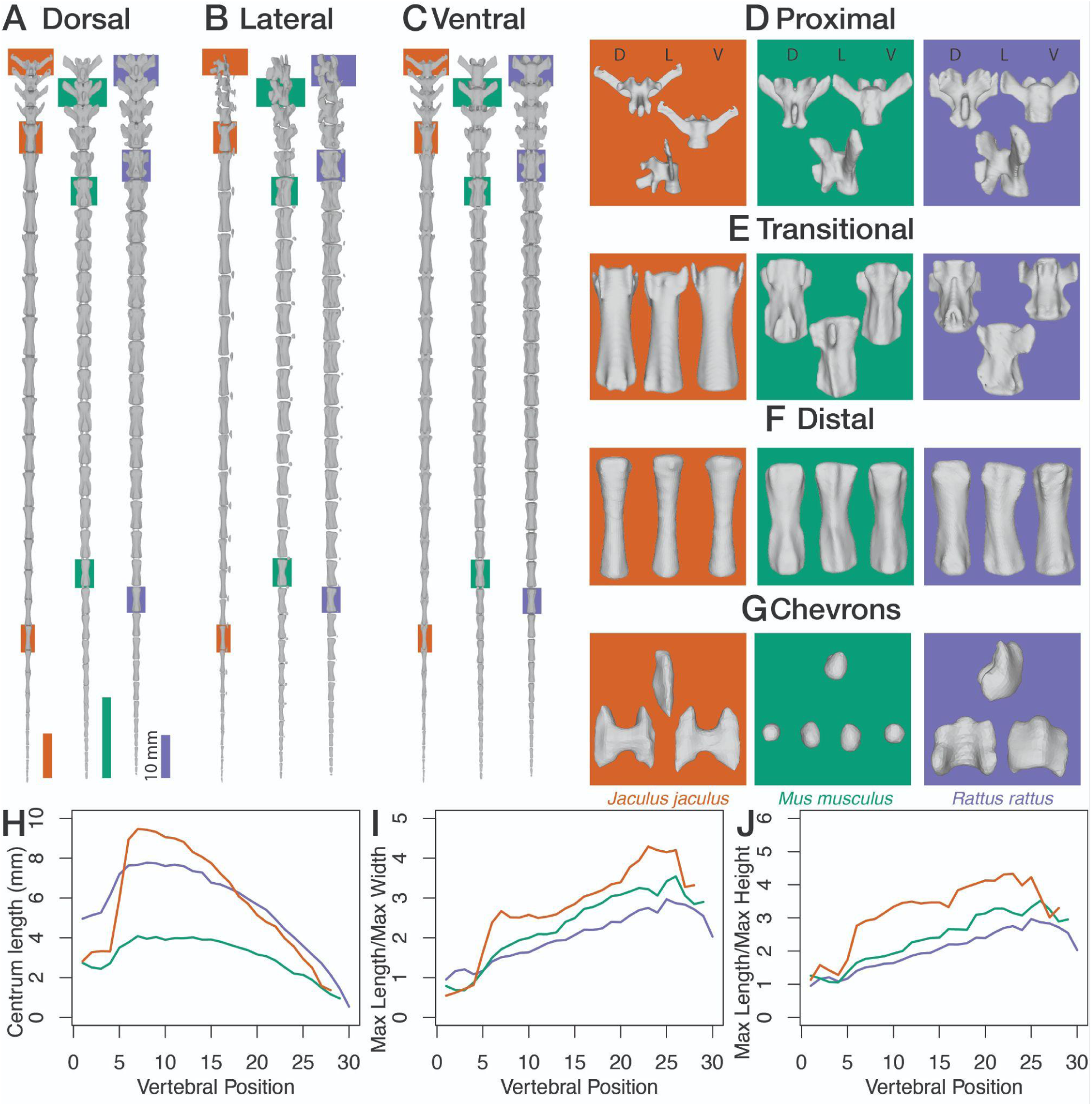
Osteological comparison across the three rodents. A) Dorsal, B) lateral, and C) ventral views of the caudal vertebral series. Colored boxes indicate which D) proximal (jerboa Cd1, mouse Cd2, rat Cd1), E) transitional (jerboa Cd5, mouse Cd6, rat Cd5), F) distal (jerboa Cd19, mouse Cd18, rat Cd19) vertebrae and G) chevrons (jerboa Cd5, mouse Cd5, rat Cd3) are shown in detail. H) Centrum length reflects the distance between joint centers of rotation, and differs from the maximum length when the transverse processes extend cranially, as in the proximal vertebrae. Jerboas have a steeper crescendo and decrescendo in centrum length than the other rodents. I) Mice and rats exhibit similar patterns of maximum length to maximum width ratio and J) maximum length to maximum height ratio, while jerboas differ in both magnitude and pattern.

The crescendo-decrescendo pattern of centrum lengths echoes the morphological regionalization of the caudal vertebrae into proximal, transitional, and distal regions which are clearly demarcated in all three rodent species we examined. The proximal vertebrae are the most sculpturally complex, as they possess neural arches topped by a spinous process on the dorsal side, a pair of laterally-projecting transverse processes, and pairs of dorsolaterally-positioned cranial and caudal articular processes or zygapophyses (*Figure 2D*). The transitional vertebrae encompass the first vertebrae with only cranial zygapophyses (i.e., no caudal zygapophyses) to the longest vertebrae (*Figure 2E*). This region is characterized by increasing simplification – loss of neural arches, reduction of zygapophyses into small rounded mammillary processes, and the shrinking and splitting of the transverse process into cranial and caudal transverse processes, connected by a lateral ridge – as well as increasing elongation to maximum centrum length. Finally, the distal vertebrae follow the decrescendo pattern of decreasing lengths and morphological simplification into rod-like forms (*Figure 2F*). We used definitions of these vertebral regions as described in rodents and primates by Ankel 1962, German 1982, Hofmann *et al*. 2021, and Organ 2010.

*Figure 2I* shows the ratio of maximum vertebral length to maximum vertebral width (*L_max_ / W_max_*), which captures the aspect ratio along the coronal plane, with higher values representing narrowing in width or increasing lateral compression of the vertebrae. All three species show a general increase in this ratio, which is to be expected as the laterally-flaring transverse processes and other processes diminish along the length of the tail. Rat caudal vertebrae showed the overall lowest length:width ratio and an abrupt decline near the tip of the tail, capturing the shortening of the terminal vertebrae. Mouse caudal vertebral length:width ratios plot just above the rat. Jerboas with their elongated vertebrae show the highest length:width ratios, with a distinct peak at the longest vertebra Cd7. However, for the first two proximal vertebrae, the jerboa show the lowest ratio values, reflecting the width contributed by the large transverse processes.

*Figure 2J* shows the ratio of maximum vertebral length to maximum vertebral height (*L_max_ / H_max_*), which captures the aspect ratio along the sagittal plane, with higher values representing narrowing in depth or increasing dorsoventral compression. All three rodent species show a similar overall trajectory in the plotted curve, with the ratio rising with vertebral position and then decreasing rapidly near the very distal end as vertebral length diminishes. Again, the mouse and rat track each other fairly closely, whereas the jerboa plots higher ratio values. Both of these aspect ratio plots – i.e., length:width and length:height – emphasize the morphology of the jerboa caudal vertebrae as elongated and slender rod-like bones.

The morphology of the caudal vertebrae is mostly similar between the mouse and rat, not only in the stocky dimensions demonstrated by comparison of aspect ratios, but also in the overall robustness of the sculptural features in the proximal and transitional vertebrae. The vertebral processes appear more prominent and larger in relation to the centrum (*Figure 2D,* purple and green). In comparison, jerboa caudal vertebral features are elegantly gracile and do not project quite as far from the centrum (*Figure 2D,* orange). For example, in the transitional vertebrae of the mouse and rat, there is a distinct, sharp lateral ridge or shelf of bone connecting the cranial and caudal transverse processes (*Figure 2E,* purple and green). This lateral ridge is barely present in the jerboa vertebrae, resulting in a smoother, lower profile (*Figure 2E,* orange). The distal vertebrae in all three species are very similar (Figure 2F). Caudal vertebrae gradually lose their morphological features distally, so the distal vertebrae take on a smooth, rod-like appearance and the tail often terminates in an irregularly-shaped nubbin of bone.

We have observed one unique morphological feature in the jerboa that occurs on Cd 6 – 10, which straddles the last few transitional and first few distal vertebrae. The cranial transverse process forms two lobes (*Figure 3A*). The primary lobe sits ventrally and, from frontal view shows a strong, hook-like ventrad curve (*Figure 3C*). The smaller auxiliary lobe sits dorsally just below the mammillary processes, giving the appearance of doubling of the mammillary processes (*Figure 3B*).

**Figure 3:**
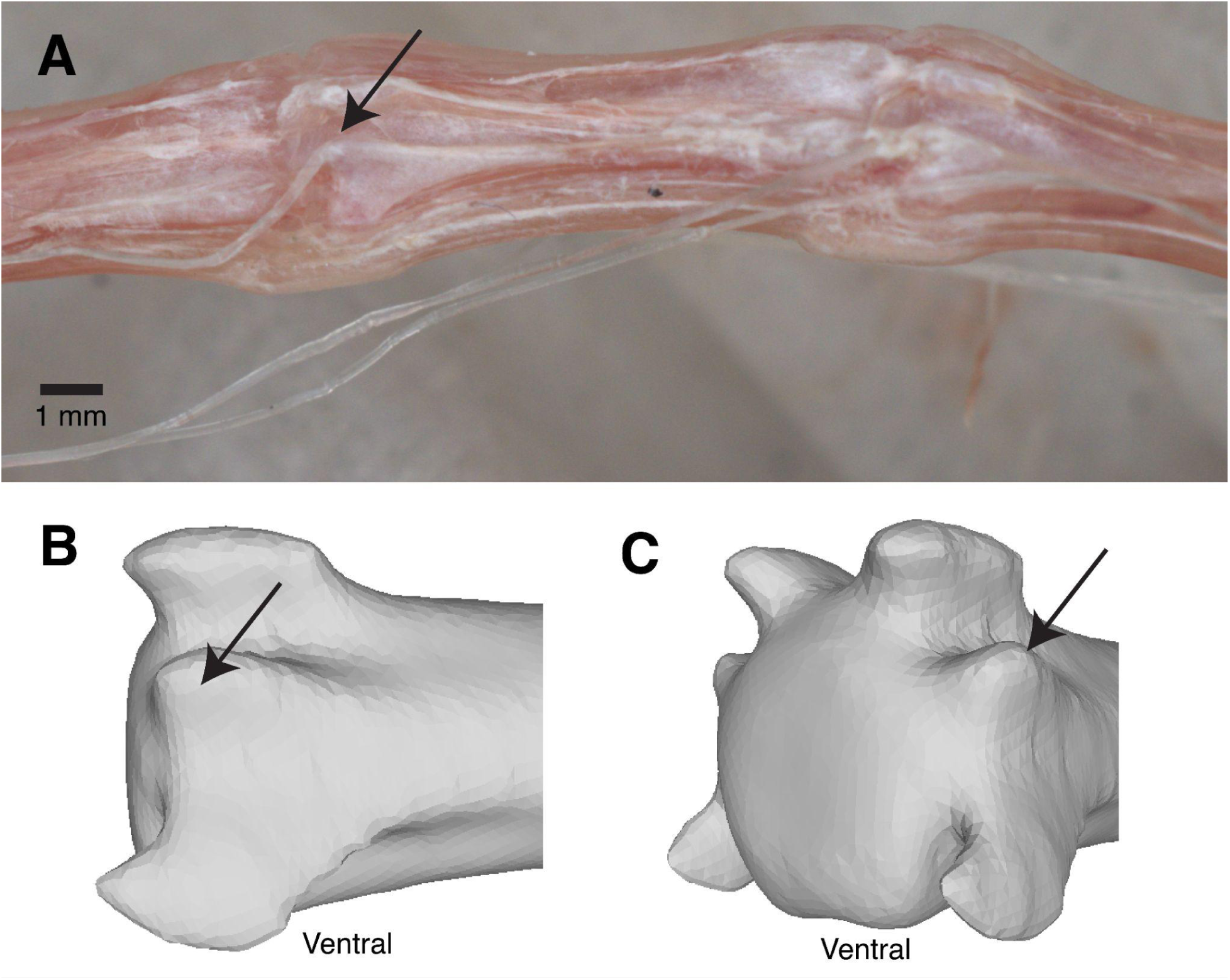
Detail of bilobed cranial transverse process, found only in jerboa. A) Photograph of dissected tail showing a tendon inserting on the dorsal lobe of the process. B) Lateral and C) oblique views of caudal vertebra 7, with arrows indicating the dorsal lobe of the transverse process. The ventral lobe of the transverse process exhibits a distinct curvature cranially

**Figure 4:**
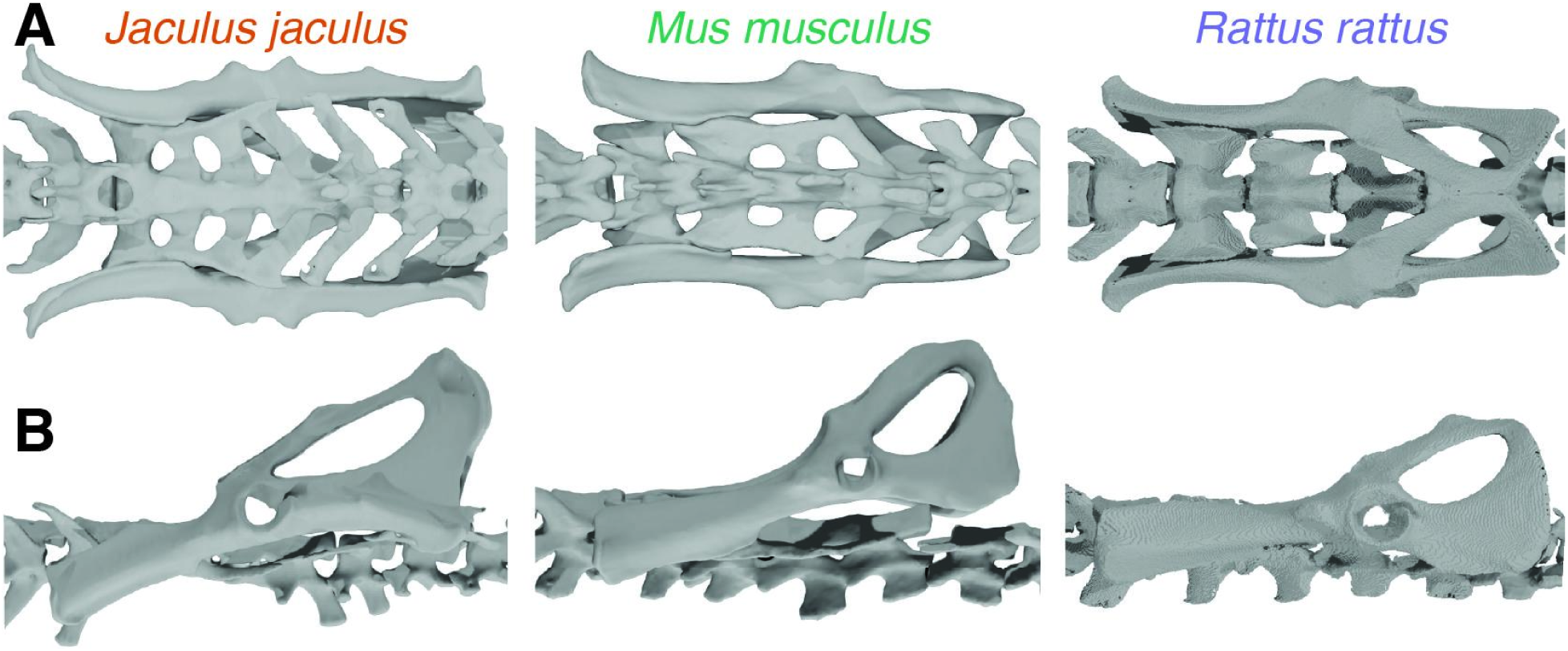
Pelvis and Sacrum details for the jerboa, mouse, and rat. A) The pelvis and sacrum are shown from a dorsal view. B) The pelvis is shown from a lateral view

### Chevrons

Chevrons, also known as haemal arches or haemal spines, are small intervertebral elements occurring along the ventral midline of the tail. These small bony elements are commonly cited as a protective structure for ventral blood vessels (Liem et al. 2001; Zavodszky and Russo 2020), but they also serve as tendon insertion sites for ventral tail muscles (Zavodszky and Russo 2020; also see section below, “Muscles”). Chevrons appear in association with the proximal caudal vertebrae and often persist throughout most of the length of the tail (*Figure 2B*). These bones become smaller and change in morphology distally, usually with the arch reducing into laterally paired bones with an irregular, pebble shape.

In the mouse, chevrons first appear between Cd 2 / 3 and the last visible chevrons occur between Cd 25 / 26. The chevrons are rounded, paired elements throughout – no arches are formed (*Figure 2G*, green). The first three proximal chevron pairs are larger in size and laterally-compressed (*Figure 2B*).

In the rat, chevrons first appear between Cd 2 / 3 and the last visible chevrons occur between Cd 22 / 23. The first three proximal chevrons are unpaired elements that form arches. The last in this series, residing between Cd 3 / 4, forms the tallest arch and has a midline keel (*Figure 2G*, purple). The remaining distal chevrons are all paired. The chevrons residing between Cd 6 / 7 to Cd 12 / 13 have a lunate shape, some of which appear disconnected due to the thinness of the bone, making pairs look like quartets (*Figure 2B*).

In the jerboa, chevrons first appear between Cd 1 / 2 and the last visible chevrons occur between Cd 22 / 23. In general, jerboa chevrons have a flat, butterfly shape (*Figure 2G*, orange) which remain as unpaired, arched elements until Cd 16 / 17; however, even the paired chevrons mostly retain their morphology and proximity to the ventral midline. The proximal chevrons have a noticeably different morphology with a dorsoventrally deep midline keel that also projects craniocaudally, giving the chevrons a pronged appearance (*Figure 2G*, orange).

### Sacrum

The sacrum is positioned over the pelvis and precedes the caudal vertebrae. Both the dorsal and ventral caudal musculature have origins and attachments on the sacrum. The sacrum consists of individual sacral vertebrae (S) that are often fused together. Connection with the pelvis is through the sacroiliac joint, where the lateral edge of the transverse processes of the sacral vertebrae contacts the interior dorsal edge of the ilium.

The mouse sacrum consists of four sacral vertebrae that show fusion at the intervertebral joints, zygapophyses, and the lateral edges of the transverse processes. The caudal end of S1 and most of S2 make up the sacroiliac joint. The sacrum has an overall rectangular shape, with relatively little change in width. The spinous processes of the neural arch for S1 – 2 are shorter, smaller, and cranially-inclined; in contrast, the spinous processes for S3 – 4 are tall, much more dorsally upright, and squared-off in appearance.

The rat sacrum consists of four sacral vertebrae that show very little evidence of fusion in the zygapophyses or transverse processes. The intervertebral joints do not show spacing indicative of the presence of intervertebral discs, but the divisions between the vertebrae are very distinct, in contrast to the mouse and jerboa, where this region is quite smoothed over. The caudal half of S1 and the cranial half of S2 make up the sacroiliac joint. Like the mouse, the sacral vertebrae are relatively uniform in width, giving the sacrum a rectangular shape. The transverse processes are broadened craniocaudally and the dorsal surface of S1 – 3 has a concave surface. The spinous processes are all oriented dorsally upright with squared-off appearance; however, the spinous process of S1 is the tallest and this height decreases in each successive vertebra.

The jerboa sacrum consists of four sacral vertebrae with fusion at the intervertebral joints, zygapophyses, and lateral edges of the transverse processes. Most of S1 and S2 make up the sacroiliac joint. The width of the sacral vertebrae varies, with some narrowing around S2 and then flaring out at S3 – 4, giving the sacrum a “waisted” or lyre-like shape. The transverse processes of S2 – 4 have a slight dorsal curl, creating a “lip.” Most the sacral vertebrae have remarkably little sculpturing, with the spinous processes and zygapophyses appearing only in very low relief. In contrast, S4 appears with large, triangular cranial zygapophyses and a slender, remarkably tall spinous process.

### Pelvis

The pelvis serves as an origination and attachment site for both dorsal and ventral caudal musculature. Dorsal musculature attaches to the dorsal edge of the ilium and the ventral musculature attaches to the interior surface of the pubis and edge of the obturator foramen.

The general shape of the pelvis is similar across all three species, though the mouse and rat share the most commonalities. These pelves have a long, narrow ilium and broad, rounded ischium/pubis with a large obturator foramen. The acetabulum is set approximately two-thirds down the length of the pelvis. The jerboa pelvis is mainly distinguished by exaggerated sculpturing of its lateral surface. The wing of the ilium has a laterally flaring curvature and the iliac crest has a pronounced lip which connects to a craniocaudally-oriented ridge that extends towards the acetabulum. The ventral edge of the ilium just cranial of the acetabulum has a large, semicircular sciatic notch, giving the base of the ilium a constricted “neck.” The ischial tuberosity is large with a laterally-projecting spine that connects to a craniocaudally-oriented ridge. These two ridges extending from the iliac crest and the ischial tuberosity form a bony shelf approximately parallel to the dorsal edge of the pelvis. The cranial edge of the pelvis in the mouse and rat is relatively flattened, but appears as a sinuous curve in the jerboa that is concave at the ischial ramus and convex at the pubis.

### Muscles

Caudal musculature is largely an extension of the axial musculature that surrounds the vertebral column and are organized into dorsal and ventral groups. The tail is also bilaterally symmetrical, which organizes the entire appendage into quadrants (i.e., left dorsal, right dorsal, left ventral, right ventral). The dorsal and ventral muscles are further divided into intrinsic and extrinsic muscles. Intrinsic muscles are small slips of muscle that connect one caudal vertebra to its adjacent neighbors. In contrast, the extrinsic muscles are large, powerful muscles anchoring on the hips and torso, sometimes communicating forces to distal caudal vertebrae via long tendons that often cross many multiple joints to reach their insertions.

The dorsal muscles – also known as the epaxial, extensor, or levator muscles – act to raise the tail dorsally and laterally. The dorsal muscles are covered by the thoracolumbar fascia over the hips and a tough connective tissue layer tightly adhering the skin to the tail.

*M. sacrocaudalis dorsalis lateralis* (SDL) is the continuation of the longissimus muscle and resides lateral to the vertebral column. This is the largest dorsal extrinsic muscle, consisting of a series of fused muscle segments and a single long tendon emerging from the end of each leaf-shaped segment. Dissections of fresh specimens allow this muscle to be excised and unfolded from their *in situ* position to reveal a spectacular segmental morphology of this muscle (*Figure 5*). As seen from this “unfolded” view, the SDL, the origin of this muscle is along its long lateral edge, which attaches to the vertebral centrum in the space between the spinous processes/zygapophyses dorsally and transverse processes ventrally. At its most cranial point, the muscle fibers of the SDL blend with the muscle fibers of the surrounding axial muscles. *In situ*, the muscle has the superficial appearance of a single fusiform belly running longitudinally down the torso since the individual segments are folded down so that the tendons are all pointed caudally. The muscle may be lightly adhered to the dorsal surface of the transverse processes of the lumbar, sacral, and proximal caudal vertebrae. The tendons insert into the cranial aspect of the mammillary processes, which are oblong processes located on the dorsocranial edge of the caudal vertebrae. Since each caudal vertebra is ostensibly attached to an SDL tendon on the left and right side, this means that individual tendons can be extremely long in order to reach the distalmost tip of the tail.

**Figure 5:**
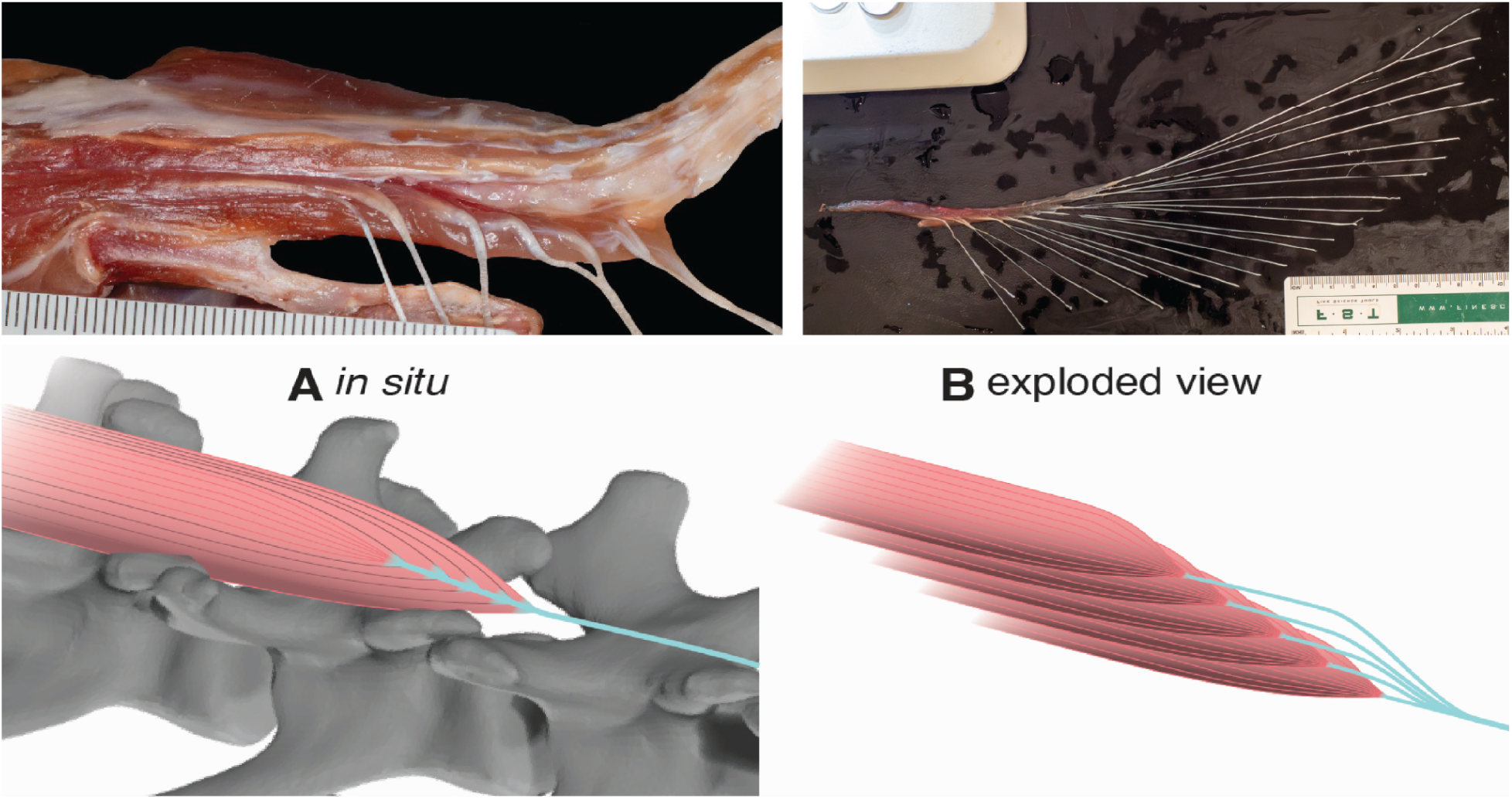
Dorsal extrinsic muscle *M. sacrocaudalis dorsalis lateralis* (SDL) in the rat in lateral view with corresponding schematic diagrams showing both (A) *in situ* position of the muscle that has been partially dissected and (B) an exploded view of the muscle after it has been excised to reveal the structure of fused muscle segments and associated tendons. Scale is in mm.

In the mouse, the SDL muscle extends from the third lumbar vertebra (L3) to Cd6 and inserts into Cd6 – 29 out of 29 total caudal vertebrae. In the rat the muscle extends from L3 to Cd7 and inserts into Cd5 – 28 out of 29 total caudal vertebrae.

In contrast, the jerboa SDL extends cranially to the penultimate thoracic vertebra (T10) to Cd5. The muscle attaches to the ventrolaterally directed transverse processes of L1 and L2 with broad tendons. A thick sheet of pelvic ligaments seems to support the SDL ventrally. The jerboa SDL appears to have a distinct medial and lateral component, with the medial component attached to the vertebral centrum as expected for the SDL. However, the lateral component attaches to the dorsal edge of the ilium and sits on the various pelvic ligaments, primarily the sacrotuberous ligament that spans from the ilium to the ischial tuberosities. In a chemically-fixed jerboa specimen, the individual muscle segments of the SDL are easily pulled apart. The more superficial muscle segments show a typical triangular or leaf shape, but deeper muscle segments begin to show a double-leafed morphology like the fletching on an arrow (*Figure 6*). The dorsal segment attaches up against the lateral wall of the spinous processes of the proximal caudal vertebrae and the ventral segment attaches on top of the transverse processes. These double-leafed muscles are limited to the lateral SDL and insert into Cd11 – 13. The tendons of the jerboa medial SDL medial component inserts into Cd6 – 10 and the lateral component inserts into Cd7 – 28 out of 28 total caudal vertebrae.

**Figure 6.**
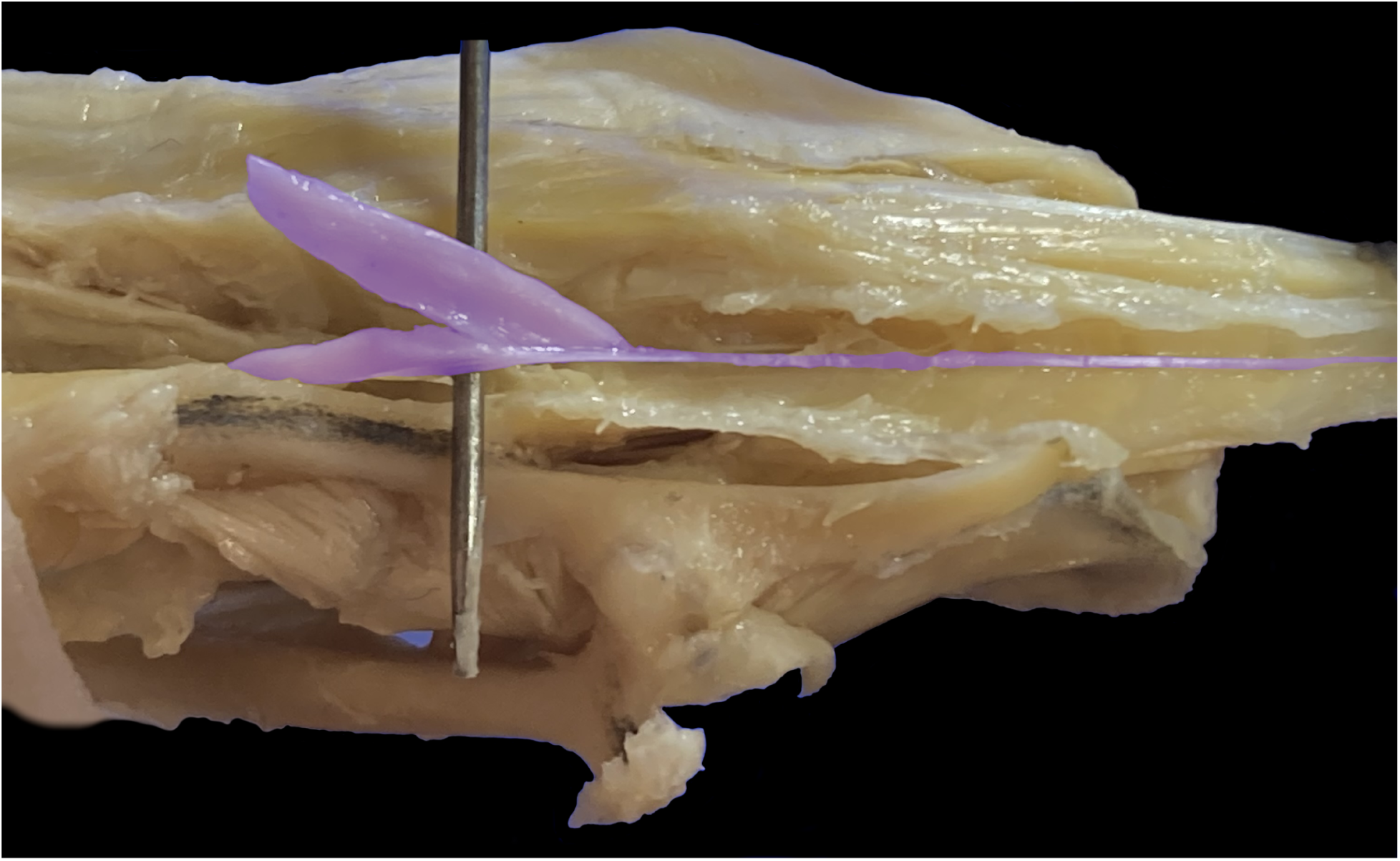
“Double-leafed” muscle segment in the jerboa, color-coded in purple.

*M. sacrocaudalis dorsalis medialis* (SDM) is the continuation of the multifidus and resides immediately parasagittal to the vertebral column and consists of short muscle segments. These segments become dorsal intrinsic muscle slips that serially repeat nearly down the entire length of the tail. All three rodent species have a similar arrangement of this muscle, but the morphology of this muscle and its tendon is unusual. A single muscle slip spans three adjacent caudal vertebrae: the muscle originates on the caudal articular process of the first vertebra, crosses over the second vertebra, and then inserts into the mammillary process of the third vertebra by sending out a short tendon that joins a long SDL tendon. This pairing of an extrinsic muscle segment with an intrinsic muscle segment which is connected by a cranially-branching tendon has been described as a “bicipital muscle” (Shinohara 1999b).These dorsal intrinsic muscles are located close to the surface of the bone and are hidden beneath the tract of dorsal tendons.

*M. intertransversarius dorsalis caudae* (ID) occurs as a fusiform or nugget-shaped muscle on the lateral surface of the base of the tail. The muscle fills the spaces between the transverse processes of the proximal tail and when freed, presents as a flattened muscle with a rounded edge extending past the dorsolateral ends of the transverse processes (*Figure 7*). This muscle is present in all three rodent species, with the most muscular development in the rat, where it extends from the pelvic ligaments at the base of the ilium to Cd6.

**Figure 7.**
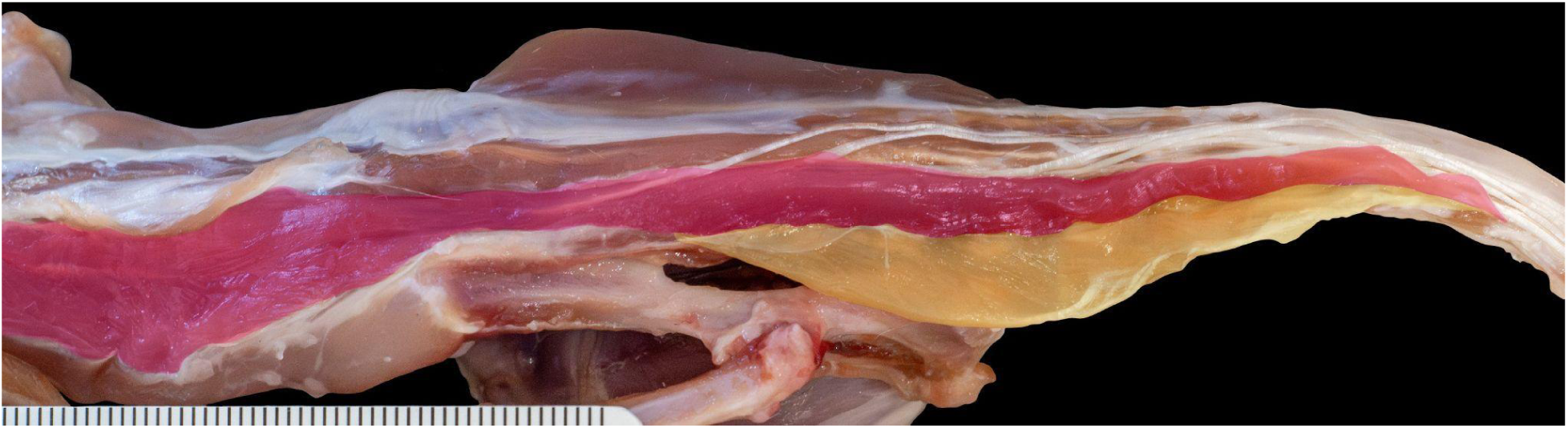
*M. intertransversarius dorsalis caudae* (ID) in the rat in left lateral view. The ID is color-coded in orange and has been pulled out laterally to show its embedding of the transverse processes and rounded shape. The SDL is color-coded pink. Scale is in mm.

The ventral muscles – also known as the hypaxial, flexor, or depressor muscles – act to lower the tail ventrally and laterally. With the exception of the *m. coccygeus* (not covered in this study), all the ventral muscles pass through the pelvic outlet alongside the rectum and urogenital tracts.

*M. levator ani* is an important muscular component of the pelvic floor which spans the pelvic outlet. These muscles originate on the interior surface of the pelvis. In all three rodent species, the levator ani is differentiated into two distinct muscles, the *m. iliocaudalis* and *m. pubocaudalis*.

The iliocaudalis originates on the interior surface of the ilium, passing over the obturator foramen, and sends out two tendons to insert either into chevrons located between vertebrae along the ventral midline or blend into the tail fascia. In the mouse, the iliocaudalis inserts between Cd5/6. In the rat, insertion was between Cd4 / 5 and Cd5 / 6. In the jerboa, this muscle also inserts tendons into the ventral midline.

The pubocaudalis is situated lateral to the iliocaudalis and originates on the interior surface of the pubis. In the mouse, this muscle inserted ventrolaterally into the tail fascia at the level of Cd3 as well as to the ventral midline. The rat pubocaudalis similarly inserted into the ventrolateral tail fascia, also at the level of Cd3. No tendons were found to arise from the jerboa pubocaudalis, though no insertions were noted.

*M. sacrocaudalis ventralis lateralis* (SVL) and *m. sacrocaudalis ventralis medialis* (SVM) occur in parallel with each other and originate parasagittally on the ventral surface of the centra of the sacral vertebrae. The SVM is usually the shorter muscle, originating caudad of the SVL. Both are multisegmented muscles with long tendons, similar in morphology to the SDL. The SVL inserts into the ventrolaterally on the cranial transverse process and the SVM inserts into the midline on the chevrons. The SVL also shows evidence of being a bicipital muscle (Shinohara 1999b), though the morphology differs slightly from the bicipital arrangement of the dorsal muscles. The ventral intrinsic muscle attaches to the caudal aspect of the caudal transverse process on the first vertebra via a tendon, extends longitudinally across the body of the second vertebra, and then attaches directly to the long tendon of the SVL which then inserts on the cranial aspect of the cranial transverse process of the third vertebra in the series. Instead of sending a short tendon to fuse with the long tendon of the SVL, the ventral intrinsic muscle appears to attach more or less directly to the long tendon. We observed one instance of a double-leafed muscle segment associated with the SVL and inserting into Cd4.

The mouse SVM muscle extends from S1 to Cd5 and its tendons insert into the ventral midline between Cd7 / 8 – 20 / 21; the SVL muscle extends from L5 to Cd3 and inserts into Cd7 - 19. In the rat, SVM muscle extends from S1 to Cd3 and inserts into the ventral midline between Cd4 / 5 – 17 / 18; the SVL muscle extends from L5 to Cd3 and inserts into Cd5 - 14. In the jerboa, the SVM muscle extends from S1 to Cd3 and inserts into the ventral midline of Cd5 / 6 – 20 / 21; the SVL muscle extends from S1 to Cd3 and inserts into Cd7 – 25. It should be noted that during dissection, many of these tendons (especially in the mouse) were broken or lost, indicating that these data may not capture the full extent of ventral insertions.

The relative contribution of dorsal versus ventral muscle is reported as the ratio of dorsal to ventral muscle weights (*Table 1*). Overall, the mouse showed greater relative ventral muscle mass and the jerboa greater relative dorsal muscle mass, while rats have an approximately equal amount of dorsal and ventral muscle. If each set of muscle weight measurements are taken as a sample, regardless of whether they came from different sides of the same animal, the mean values of the dorsal:ventral ratios are: mouse = 0.90 (n = 5), rat = 1.01 (n = 2), and jerboa = 1.18 (n = 2). Mean values of dorsal:ventral ratios if values were averaged per individual are: mouse = 0.86 (n = 3), rat = 1.01 (n = 1), and jerboa = 1.18 (n = 1). We collected weight measurements from both wet and dried muscles: mouse wet = 0.92 (n = 2) vs. dry = 0.84 (n = 3); rat wet = 0.99 (n = 1) vs. dry = 1.03 (n = 2); jerboa wet = 1.07 (n = 1) vs. dry = 1.29 (n = 1). Though these values shift depending on weight collection method and binning of samples, the relationship of the mouse, rat, and jerboa dorsal:ventral ratios remains the same.

**Table 1.** Dorsal versus ventral muscle weights.

| SPECIES | SPECIMEN ID | SEX | WET or DRY? | DORSAL WT<br>MEAN (g) | VENTRAL WT<br>MEAN (g) | DORSAL:<br>VENTRAL |
| --- | --- | --- | --- | --- | --- | --- |
| *Mus musculus* | 2024.12.17 | ? | Dry | 0.022 | 0.027 | **0.81** |
| *Mus musculus* | 2024.12.26 | ? | Wet<br>(right side) | 0.125 | 0.152 | **0.82** |
| *Mus musculus* | " | " | Wet<br>(left side) | 0.126 | 0.124 | **1.02** |
| *Mus musculus* | 2025.10.26 | M | Dry<br>(right side) | 0.037 | 0.041 | **0.90** |
| " | " | " | Dry<br>(left side) | 0.040 | 0.050 | **0.80** |
| *Rattus norvegicus* | 2024.12.20 | ? | Wet<br>(right side) | 2.631 | 2.661 | **0.99** |
| " | " | " | Dry<br>(left side) | 0.561 | 0.546 | **1.03** |
| *Jaculus jaculus* | JJ0998 | F | Wet | 0.306 | 0.287 | **1.07** |
| " | " | " | Dry | 0.085 | 0.066 | **1.29** |

### Tendons

When the skin of the tail is removed, the most striking feature are the bundles of tightly packed tendons at each quadrant, running down the entire length of the tail. These extraordinarily long tendons emerge from the SDL and SVL muscles, individually inserting into the bones of the tail, even crossing many multiple joints to reach the terminal vertebra at the distalmost tip. The first few proximal vertebrae over the hips and at the base of the tail do not have tendon insertions, but instead serve as a sort of staging area for the extrinsic caudal muscles. However, each remaining caudal vertebra is ostensibly attached to six tendons: left and right SDL muscle tendons inserting into the dorsolateral mammillary processes, left and right SVL muscle tendons inserting into the ventrolateral cranial transverse processes, and the left and right SVM muscle tendons inserting onto either side of the chevrons along the ventral midline. Multiplying six tendons per vertebra in a tail consisting of over 20 caudal vertebrae results in a staggering number of individual tendons. “Cable management” of the tail tendons is critical to maintaining the most direct line of action with the muscles in a flexible appendage while avoiding tangling or interference. This is primarily achieved through connective tissue. The tail itself is tightly covered in the caudal fascia, but each tendon is individually wrapped in a sheath of connective tissue and tendons in a tract are bundled together in a fascicle which adheres to the bone. The tendons in a tract run in parallel and stacked on top of each other. Typically, the tendons are organized so that the ones inserting more proximally are located superficially and the tendons inserting distally are deeper in the fascicle.

We have observed some unusual tendon morphologies in the tail, viz., two forms of tendon branching. The first form of branching has already been discussed above in the description of the bicipital muscles. The long tendons from a muscle segment of the SDL will show a short, cranially-branching tendon attached to a small intrinsic muscle slip, thus pairing between an extrinsic and intrinsic muscle segment together. The second form of branching occurs in the long tendons of the SDL and SVL, where a single tendon emerging from a single muscle segment will distally split into two or as many as six branches in the jerboa, each one inserting into different adjacent vertebrae (*Figure 8*). In addition, the ventral caudal musculature has a tendency to insert into the caudal fascia via tendons that are likewise flat and sheetlike.

**Figure 8.**
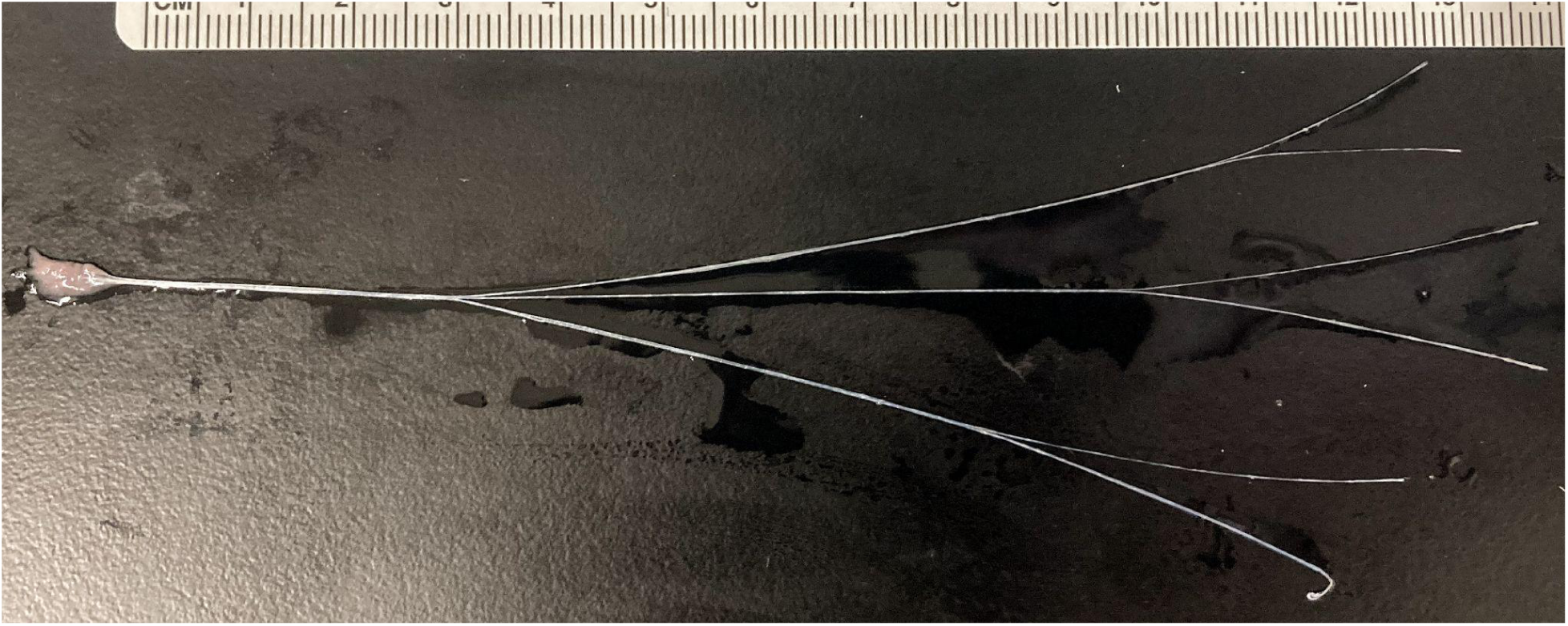
Jerboa muscle segment showing branching in a long tendon. Scale is in mm.

There are few outstanding differences in the tendons of the mouse, rat, and jerboa. The technical difficulty of tendon microdissection in this study has resulted in some regions of missing data, but there is sufficient evidence that when all tendon insertions are mapped out on the caudal vertebrae, there appears to be considerable individual variation. Not all caudal vertebrae have six tendon attachments, having more or less than the expected number (*Figure 9*). One species-specific variation is in the jerboa, which shows supernumerary dorsal tendon attachments around Cd6 – 10, a region corresponding to the presence of a bi-lobed cranial transverse process which acts as an additional tendon attachment site for the SDL.

**Figure 9:**
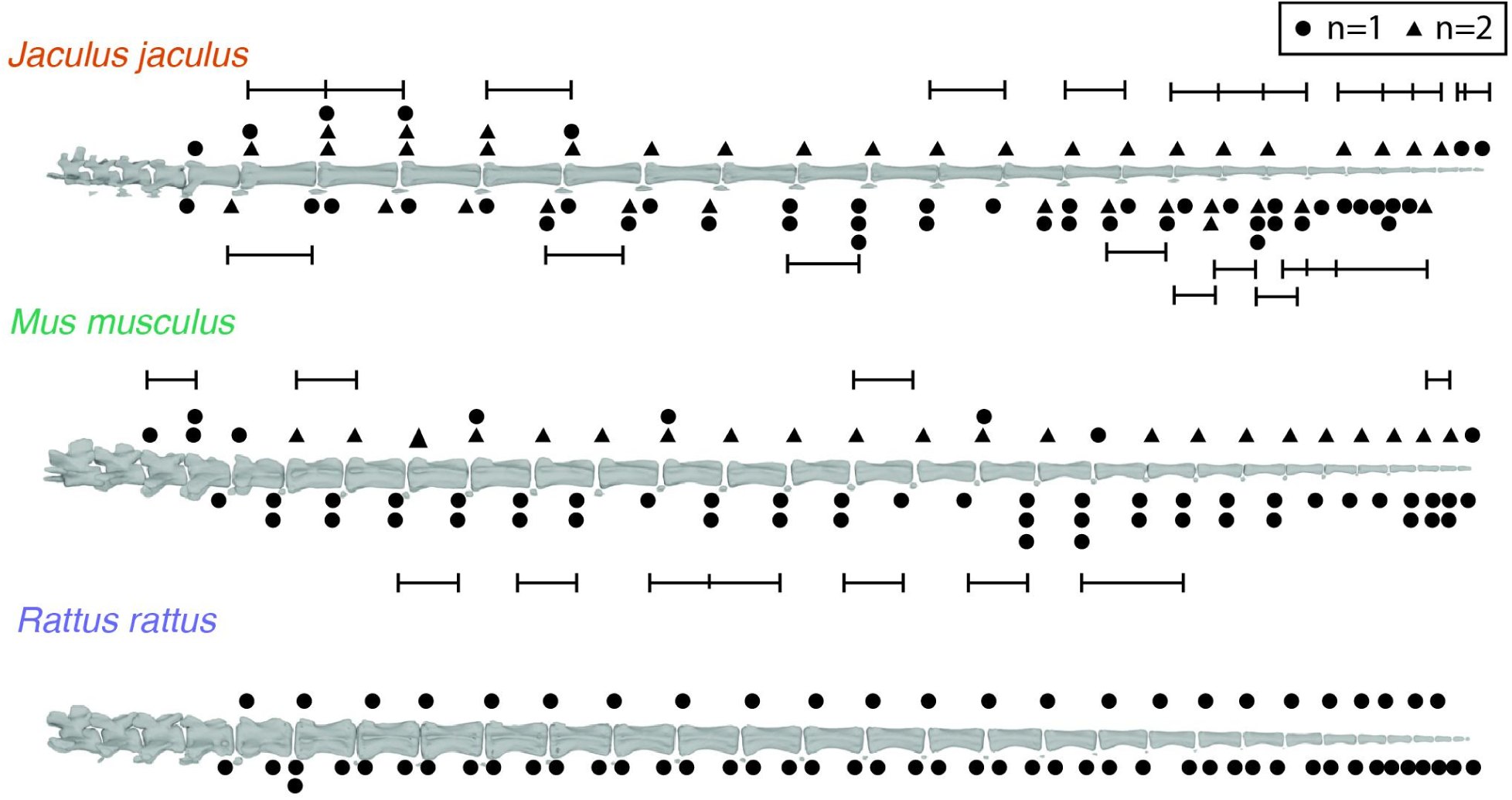
Diagram of tendon insertions. Circles represent insertions found in one specimen. Triangles represent insertions found in two specimens. Brackets indicate insertions that are connected via branched tendons.

## DISCUSSION

### Functional interpretation of anatomy

Many aspects of the structure, organization, and regionalization of the tail are conserved across our comparative sample of three rodent species. However, our focal species, the lesser Egyptian jerboa (*Jaculus jaculus*), demonstrates some interesting derived features that may have some functional relevance to the proposed role of the tail as a rapidly moving inertial appendage used to facilitate quick changes in body orientation during escape flights. Jerboas have strikingly long tails, approximately 1.69 times their body length, compared to the mouse at 0.96 and rat at 1.05, from our sample. Despite these radically different lengths, all three rodents share similar numbers of caudal vertebrae. The jerboa achieves its remarkable tail length through the elongation of individual vertebrae, which takes place during endochondral bone development by increasing both the number and size of chondrocytes that build the cartilaginous scaffold for bone ossification (Weber et al. 2025). Longer tails have been experimentally determined with animal trials and tailed robots to be more effective inertial appendages than shorter ones (Libby et al. 2012). Furthermore, the vertebrae in the tail do not change length uniformly. All three rodents show a crescendo-decrescendo pattern of vertebral centrum length change, but jerboas showed the steepest curve with the length of each vertebra increasing rapidly to its maximum length in Cd7 before decreasing again. A computational model of a simulated linked tail attached to a body showed that given a task to reorient the body along a predetermined trajectory using tail movements, the tail with link lengths most closely resembling the jerboa showed the lowest tracking error compared with tails with uniform link length or even link lengths with a much less pronounced crescendo-decrescendo, similar to those that would be found in the mouse and rat (Fu et al. 2025).

The morphology of the jerboa caudal vertebrae is surprisingly gracile, not only due to the elongation of the bones, but also in the comparatively delicate and low-profile shape of the processes. This elegant sculpting continues to the sacrum and pelvis, where the extrinsic muscles reside as the primary drivers of tail movement, but these bones show features for large muscle attachments. For example, the sacrum is broadened distally with long, dorsally concave transverse processes to support the placement and attachment of the sacrocaudalis dorsalis lateralis (SDL) muscle. The lack of bulky, robust processes for tendon and intrinsic muscle attachment on the caudal vertebrae themselves may be weight-saving for an appendage that has evolved to whip around at high speeds. However, there are potential alternative strategies for supporting powerful muscular forces such as retention of the chevrons’ midline-straddling morphology to the distal end of the tail when mouse and rat chevrons have diminished into small, paired pebbles of bone on either side of the ventral midline, thus possibly providing more stable support to the sacrocaudalis ventralis medialis (SVM) tendons inserting there. The bi-lobed cranial transverse process is a novel feature found in jerboas and is discussed at length below.

We observed some changes in jerboa muscle morphology, especially the SDL, which is the principal dorsal extensor. Unlike the SDL in the mouse and rat which have their cranial origin on the third lumbar vertebra, the jerboa SDL originates much far forward near the penultimate thoracic vertebra, T10. This muscle, which typically attaches along its long medial edge on the vertebral centra in the slot between the spinous process/zygapophyses dorsally and transverse processes ventrally, the jerboa SDL differentiates into a lateral and medial component. The medial component continues to attach to the vertebral centra, but the lateral component attaches to the dorsal edge of the ilium, which we hypothesize may provide not only an additional skeletal anchor, but perhaps also a more stable one compared to the vertebral column. We see another doubling of muscular attachment in the deeper muscle segments of the SDL and SVL, where two muscle segments braced on different skeletal elements are found on a single tendon. Finally, an assessment of the ratio of dorsal to ventral muscle weights show that the jerboa (1.18) has greater dorsal muscle mass compared to the mouse (0.90) and rat (1.01). This additional investment in dorsal muscle mass, supported by additional skeletal attachments, may be necessary for the jerboa to lift and move its substantially longer tail.

#### Novel anatomical feature: bi-lobed cranial transverse processes

We noted an unusual morphology in caudal vertebrae Cd 6 – 10 in the jerboa: bi-lobed cranial transverse processes, with a large, ventrally-curved ventral lobe and a smaller, dorsolaterally projecting dorsal lobe (*Figure 3B, C*). This series of vertebrae consists of the last two transitional vertebrae, including the longest vertebra, and the first three distal vertebrae. A cursory examination for the presence of this morphology in a small sample of other rodent species (for details, please see Supplemental Materials, *Table 4*) found bi-lobed cranial transverse processes are present only in other members of the Family Dipodidae (*Allactaga severtsovi sungorus* and *Salpingotus crassicauda*). However, members of sister clades within Superfamily Dipodoidea do not share this morphology (*Napaeozapus insignis frutectanus*, *Zapus hudsonius*). Other bipedal hopping rodents also lack this feature (*Notomys alexis, Dipodomys merriami merriami*). This information suggests that the bi-lobed cranial transverse process may be a synapomorphy for Dipodidae, but because we do not have data on whether other jerboa species use their tails in a similar manner to *J. jaculus,* it remains uncertain whether this is a mere phylogenetic signal or there is functional relevance.

In our dissections, we observed the cranial aspect of the dorsal lobe as a site for tendon attachments from the SDL (sacrocaudalis dorsalis lateralis FIGURE 3A). Instead of the expected single tendon per dorsal left/right side inserting into the mammillary process, jerboa tendon maps show an additional one or two supernumerary tendon attachments for vertebrae in the region that overlap with the incidence of these bi-lobed cranial transverse processes. The additional tendon attachment site that the dorsal lobe provides suggests the importance of these caudal vertebrae as a site where muscular force and/or control needs to be concentrated during the rapid whipping movements of the tail when it is functioning as an inertial appendage. In contrast, the potential functional role of the ventral lobe is much less apparent. The ventral intrinsic muscles were observed passing underneath the ventral lobe, but it is unclear whether these small muscles benefit from being sheltered by this extension of bone or if the bone is presenting an extra attachment surface. Broadly, it is possible that the ventrally-directed curvature or hook of the ventral lobe acts to help reinforce the position of the ventral tendon tract in a high force load area since any displacement or rupture of individual tendons from their connective tissue sheath would likely be incapacitating. Research on primates proposes that the longest caudal vertebra will experience the highest level of bending and torsion (Organ 2010). Since the presence of bi-lobed cranial transverse processes encompasses the longest caudal vertebra and adjacent bones which are similarly elongated, it is possible that this feature may serve a protective role.

#### Novel anatomical feature: branching tendons

Another striking feature we found was two types of tendon branching in the tail, which was found in all three species we examined. The first type of branching connects together muscle segments in the sacrocaudalis dorsalis or sacrocaudalis ventralis to an intrinsic muscle slip through a short, cranially-directed forking of the tendon just before its insertion site, thus forming what Shinohara (1999b) calls a “bicipital muscle,” referencing the presence of both an extrinsic and intrinsic muscle head. Though there is no explicit description or clear visual depiction of branching, Shinohara does mention in passing of how the tendons from the two muscles “join.…and are inserted into the cranial articular process.…” Whether the tendons are splitting into branches or joining together is a developmental question that is not attended to in our current study. As to the functional implications, Shinohara speculates that the contraction of the short intrinsic muscles might not only provide regional movements, but also act to pull the long extrinsic muscle tendon medially to enhance the efficacy of action. Without knowing whether both muscles in the bicipital pair contract simultaneously or in some other muscle activation pattern, it is difficult to determine how the extrinsic and intrinsic muscle pairs work together. However, we posit a similar hypothesis to Shinohara (1999b) and Hori (2011) that the bicipital arrangement may allow for the exertion of localized control of joint stiffness, allowing for more precise tail movements.

The second form of branching we found was in the long tendons coming from the sacrocaudalis dorsalis and sacrocaudalis ventralis muscles. This morphology arises from what appears to be a single muscle segment sending out a single tendon which then branches distally and each of those branches typically insert into different, but adjacent caudal vertebrae. The distribution of this sort of tendon branching is much more variable both between and within species. For dissection specimens with tendon maps, no such branching was found in the rat (n = 1), but branching was found in one mouse (n = 2) and in all jerboas (n = 3). In the jerboas, the number of branched tendons and the position of vertebrae receiving attachments from branched tendons was highly variable. Therefore, it is challenging to assign a functional significance for branching in these tendons. An alternative hypothesis is that branching in these long tendons is a developmental artifact where neighboring muscle segments and their tendons were not completely split into separate parts during morphogenesis. We have observed different degrees of splitting with some putatively branching tendons actually arising from closely adhered muscle segments, as well as spatial groupings of two or more distinct tendons arising as a cluster from adjacent muscle segments. Genuine instances of branching occur when branches are pulled apart and eventually show signs of fraying from a blending of collagen fibers. However, what causes such irregularities in the development in what is ostensibly a regular, metameric anatomical system is unresolved.

#### Limitations and considerations of this comparative study

This study describes the musculoskeletal anatomy of the tail in three species of rodent with the intent of identifying a correlation between form and function, especially in relation to the role of the tail in locomotion. In addition to the substantial technical difficulty of tail muscle and tendon microdissection leading to incomplete and misinterpreted anatomical data, any comparative study faces inherent limitations in sampling and conceptual framing that obscure or occlude these form-function signals. We present three points of consideration that acknowledge these limitations and shift our perspective towards a potentially more complex, holistic understanding of tails.

First, our gross morphological description critically included tail to body length ratios. We reported ratios for both lab-bred and housed animals that we measured directly and ratios calculated from measurements taken from wild animals deposited in natural history collections and made freely available online via GBIF. While the average tail to body length ratios for mice and jerboas differ little between captive and wild populations (0.96 vs. 0.96 and 1.69 vs. 1.66, respectively), we find a notable difference between laboratory (1.05) and wild rats (0.84). In rats, this ratio can shift with strain, age, and sex, with females tending towards having slightly longer tails (Bao 1995). The salvaged rat carcass we used for our dissection is likely a Sprague Dawley, an outbred strain of albino rat popularly used in laboratory studies and morphologically characterized by having a tail length exceeding body length (Janvier Labs, n.d.), which likely accounts for the discrepancies in proportions. Strain-specific phenotypes are a significant factor, but not the only drivers of tail morphology in laboratory animals. For instance, it has long been documented that rearing mice in cold versus warm environments results in longer tails in the latter (Barnett et al. 1975; Freymann et al. 2017) or that mice limited to climbing locomotion in simulated arboreal housing increased the length of the transverse processes in their transitional caudal vertebrae (Byron et al. 2011). This serves to highlight how, despite the goal of describing tail anatomy in its most representative and neutral state, the process of establishing clear form-function relationships can be confounded by the artificiality of captivity in our study subjects in both recognized and unrecognized ways.

Secondly, we were motivated to understand the anatomical foundations for the jerboa’s tail functions, but to determine the outstanding characteristics unique to the jerboa requires careful selection of species with which to make a comparison. We chose the laboratory mouse and rat, not only because they are readily available with an existing body of literature on tail anatomy (Brink and Pfaff 1980; Hori et al. 2011; Shinohara 1999b), but also because those species are frequently considered the standard generalist quadrupedal in many comparative studies (e.g., Charles et al. 2016; Polly 2007; Schmidt and Fischer 2011; Sears et al. 2006; Wright et al. 2022), a sort of “null” locomotory hypothesis against which the derivation of characters is assessed. But it should be noted that both mice and rats have been observed using their tails in specialized ways: when placed on unstable narrow beams or rods, mice and rats will swing their tails around as an active counterbalance and even wrap their tails around the rod to prevent falling and help righting themselves when upside down (Buck et al. 1925; Hori et al. 2011). While evidence of such behavior should call into question the extent to which laboratory mice and rats represent a truly generic quadruped for comparative studies, the semi-prehensile use of the tail during climbing is an apt comparison, as the birch mice (Sminthidae) are close extant relatives of jerboas that are phylogenetically placed at the base of Dipodoidea and show quadrupedal, semi-arboreal locomotion, which most probably represents the plesiomorphic state for this group (Moore et al. 2015).

Lastly, our interest in the jerboa was inspired by the potential contribution of their long tails to their highly specialized modes of bipedal locomotion. While locomotion is hypothesized to be the primary driver for changes in tail morphology and anatomy, it remains important to take into consideration other factors for selection. For instance, we find evidence that jerboas have greater dorsal versus ventral caudal muscle mass, which would support the notion that greater muscular contribution to tail extension in jerboas may be part of an adaptive suite of specializations for ricochetal bipedal locomotion that incorporate rapid, wide-range motions of the tail to aid in quick directional changes. However, our observations of captive jerboas show that the tail is erected to a near-vertical position by the pursuing male during courtship (*Supplemental Figure 1*). This would suggest potential sexual dimorphism in muscle distribution, but our current sample of specimens is inadequate to test this. Future studies will need to address the influence of other functional roles and evolutionary drivers for shaping the anatomy of the tail in this species.

We are only beginning to understand the function and biomechanics of mammalian tails, especially in a comparative framework appropriate for understanding evolutionary patterns, form-function relationships, and applications to bioinspired engineering and design. The work presented here integrates aspects of osteology with soft-tissue muscle and tendon anatomy across three species of rodents to help establish a solid foundation from which such studies and applications can emerge. We have contributed comparative musculoskeletal anatomical descriptions of the mouse, rat, and jerboa tail while identifying some key features in the jerboa such as elongated caudal vertebrae, potentially novel bi-lobed cranial transverse processes serving as an auxiliary dorsal tendon attachment site and reinforcement of ventral tendon position, and greater dorsal versus ventral muscle mass. These features support the hypothesis that the jerboa tail is adapted for fast, powerful movements such as those observed during ricochetal locomotion. Caudal muscles and tendons are an intricate and complex anatomical system that remain a rich source of scientific inquiry across many disciplinary perspectives.

## Supporting information

Miyamae & Moore_Tail Anatomy of the Jerboa_Supplemental

## ACKNOWLEDGEMENTS

JAM and TYM were supported by University of Michigan Robotics and NSF IntBIO Grant 2500297. Specimens used for this study were obtained through the generosity of Matthew Boulanger, Sarah Nowlan, and other members of the Animal Care and Use Program training core at the University of Michigan at Ann Arbor; Kimberly Cooper, Ceri Weber, and Swithin Razu at the University of San Diego. MicroCT scans for this study were made available thanks to MorphoSource (morphosource.org). TYM thanks Daniel Johnson for assistance with Blender.

## Notes

### Competing Interest Statement

The authors have declared no competing interest.

