## Supplementary material for "Specializations in Tail Anatomy of the Lesser Egyptian Jerboa (*Jaculus jaculus*) Compared with the Mouse and Rat": Miyamae & Moore_Tail Anatomy of the Jerboa_Supplemental

SUPPLEMENTAL MATERIALS

*Table 1*. Tail:body length for specimens measured in this study.

| SPECIES | SPECIMEN ID | SEX | TAIL LENGTH (mm) | BODY LENGTH (mm) | TAIL : BODY |
|---|---|---|---|---|---|
| *Mus musculus* | 2024.08.20 | ? | 84 | 96 | 0.88 |
| *Mus musculus* | 2024.12.26 | ? | 86 | 93 | 0.92 |
| *Mus musculus* | 2025.09.01 | F | 110 | 102 | 1.08 |
| *Mus musculus* | 2025.10.26 | M | 91 | 97 | 0.94 |
| *Rattus norvegicus* | 2024.12.20 | ? | 239 | 228 | 1.05 |
| *Jaculus jaculus* | UCSD JJ0998 | F | 177.2 | 106.1 | 1.67 |
| *Jaculus jaculus* | UCSD JJ0360 | M | 179 | 104 | 1.72 |
| *Jaculus jaculus* | UCSD JJCWAM 101922125 | M | 167.93 | 99.47 | 1.69 |

*Table 2*. Tail:body length data for museum specimens reported in GBIF.

| SPECIES | SPECIMEN ID | SEX | TAIL LENGTH (mm) | BODY LENGTH (mm) | TAIL : BODY |
|---|---|---|---|---|---|
| *Mus musculus* | AMNH M-270961 | F | 76 | 85 | 0.89 |
| *Mus musculus* | UCONN 11347 | F | 65 | 67 | 0.97 |
| *Mus musculus* | EMU M 589 | M | 82 | 77 | 1.06 |
| *Mus musculus* | UNSM 14480 | M | 71 | 76 | 0.93 |
| *Mus musculus* | MCZ Mamm:63034 | M | 71 | 79 | 0.90 |
| *Mus musculus* | CUMV 7795 | M | 66 | 87 | 0.76 |
| *Mus musculus* | UWBM Mamm:61749 | F | 72 | 67 | 1.07 |
| *Mus musculus* | UWBM Mamm:9217 | F | 91 | 91 | 1 |
| *Mus musculus* | UWBM Mamm:37048 | F | 82 | 72 | 1.14 |
| *Mus musculus* | ZMBN:OM:007773 | M | 67 | 79 | 0.85 |
| *Rattus norvegicus* | UCONN 14749 | F | 114 | 137 | 0.83 |
| *Rattus norvegicus* | CRCM Mamm:10632 | F | 168 | 212 | 0.79 |
| *Rattus norvegicus* | NML 1986.102.11 | F | 161 | 193 | 0.83 |
| *Rattus norvegicus* | MSB Mamm:291707 | F | 130 | 165 | 0.79 |
| *Rattus norvegicus* | UAM Mamm:101039 | M | 215 | 206 | 1.04 |

| | | | | | |
|---|---|---|---|---|---|
| *Rattus norvegicus* | UAM Mamm:111949 | M | 198 | 249 | 0.80 |
| *Rattus norvegicus* | UAM Mamm:154123 | M | 135 | 176 | 0.77 |
| *Rattus norvegicus* | KSTC Mammals:AA0031 | M | 150 | 163 | 0.92 |
| *Rattus norvegicus* | UCONN 17081 | M | 85 | 105 | 0.81 |
| *Rattus norvegicus* | CUMV 7756 | F | 150 | 191 | 0.78 |
| *Jaculus jaculus* | CUMV 8861 | M | 183 | 110 | 1.66 |
| *Jaculus jaculus* | CUMV 13381 | F | 186 | 109 | 1.71 |
| *Jaculus jaculus* | CUMV 11675 | M | 161 | 92 | 1.75 |
| *Jaculus jaculus* | CUMV 13384 | M | 169 | 101 | 1.67 |
| *Jaculus jaculus* | CUMV 13383 | M | 178 | 108 | 1.65 |
| *Jaculus jaculus* | CUMV 13382 | M | 189 | 109 | 1.73 |
| *Jaculus jaculus* | CUMV 13385 | M | 183 | 118 | 1.55 |
| *Jaculus jaculus* | UCM Mamm:14759 | M | 178 | 116 | 1.53 |

Institutional abbreviations: **AMNH**, American Museum of Natural History; **CRCM**, Washington State University Charles R. Conner Museum; **CUMV**, Cornell University Museum of Vertebrates; **EMU**, Eastern Michigan University T.L. Hankinson Vertebrate Museum; **KSTC**, Schmidt Museum of Natural History; **MCZ**, Harvard University Museum of Cmoparative Zoology; **MSB**, Museum of Southwestern Biology; **NML**, National Museums Liverpool; **UAM**, University of Alaska Museum of the North; **UCM**, University of Colorado Museum of Natural History; **UCONN**, University of Connecticut Biodiversity Research Collections; **UNSM**, University of Nebraska State Museum; **UWBM**, University of Washington Burke Museum; **ZMBN**, The Mammal Collection of the University Museum of Bergen

*Table 3*. Vertebral counts.

| SPECIES | SPECIMEN ID | SPECIMEN SOURCE | SEX | # of CAUDAL VERTEBRAE |
|---|---|---|---|---|
| *Mus musculus* | 2024.08.20 | Dissection | ? | 29 |
| *Mus musculus* | 2024.12.17 | Dissection | ? | 30 |
| *Mus musculus* | 2025.09.01 | Dissection | F | 32 |
| *Mus musculus* | TMM.M.8671 | MicroCT scan (MorphoSource media ID 000606487) | ? | 29 |
| *Mus musculus* | UMZC.E.2204 | MicroCT scan (MorphoSource media ID 000602037) | ? | 32 |
| *Rattus norvegicus* | 2024.12.20 | Dissection | ? | 29 |
| *Rattus norvegicus* | UMMZ Mamm 167022 | MicroCT scan (MorphoSource media ID 000082216) | F | 30 |
| *Jaculus jaculus* | 2022.07.22 | Dissection | ? | 27 |
| *Jaculus jaculus* | 2023.07.08 | Dissection | F | 28 |
| *Jaculus jaculus* | 2024.04.03 | Dissection | M | 28 |

| | | | | |
|---|---|---|---|---|
| *Jaculus jaculus* | 2025.01.26 | Dissection | F | 28 |
| *Jaculus jaculus* | UMMZ Mamm 120204 | MicroCT scan (MorphoSource media ID 000064399) | ? | 28 |

Institutional abbreviations: **TMM**, University of Texas Vertebrate Paleontology Collections; **UMMZ**, University of Michigan Museum of Zoology; **UMZC**, University Museum of Zoology in Cambridge

*Table 4.* Rodent species examined for the presence of bi-lobed cranial transverse processes.

| SPECIES | FAMILY, SUPERFAMILY | SPECIMEN ID | BILOBED CRANIAL TRANSVERSE PROCESSES? |
|---|---|---|---|
| *Allactaga severtsovi sungorus* | Dipodidae, Dipodoidea | UMMZ Mamm 123053 (MorphoSource media ID 000570333) | yes |
| *Jaculus jaculus* | Dipodidae, Dipodoidea | UMMZ Mamm 120204 (MorphoSource media ID 000064399) | yes |
| *Salpingotus crassicauda* | Dipodidae, Dipodoidea | UMMZ Mamm 120123 (MorphoSource media ID 000068587) | yes |
| *Napaeozapus insignis frutectanus* | Zapodidae, Dipodoidea | UMMZ Mamm 126314 (MorphoSource media ID 000056072) | no |
| *Zapus hudsonius* | Zapodidae, Dipodoidea | YPM MAM 005659 (MorphoSource media ID 000064824) | no |
| *Notomys alexis* | Muridae, Muroidea | SAMA M24165 (MorphoSource media ID 000500070) | no |
| *Dipodomys merriami merriami* | Heteromyidae, Geomyoidea | UMMZ Mamm 120461 (MorphoSource media ID 000569151) | no |

Institutional abbreviations: **UMMZ**, University of Michigan Museum of Zoology; **SAMA**, South Australian Museum Australia; **YPM**, Yale Peabody Museum of Natural History

*Figure 1*. Still images from video footage of male-female pair of captive jerboas during courtship, showing the pursuant male with tail held vertically aloft.

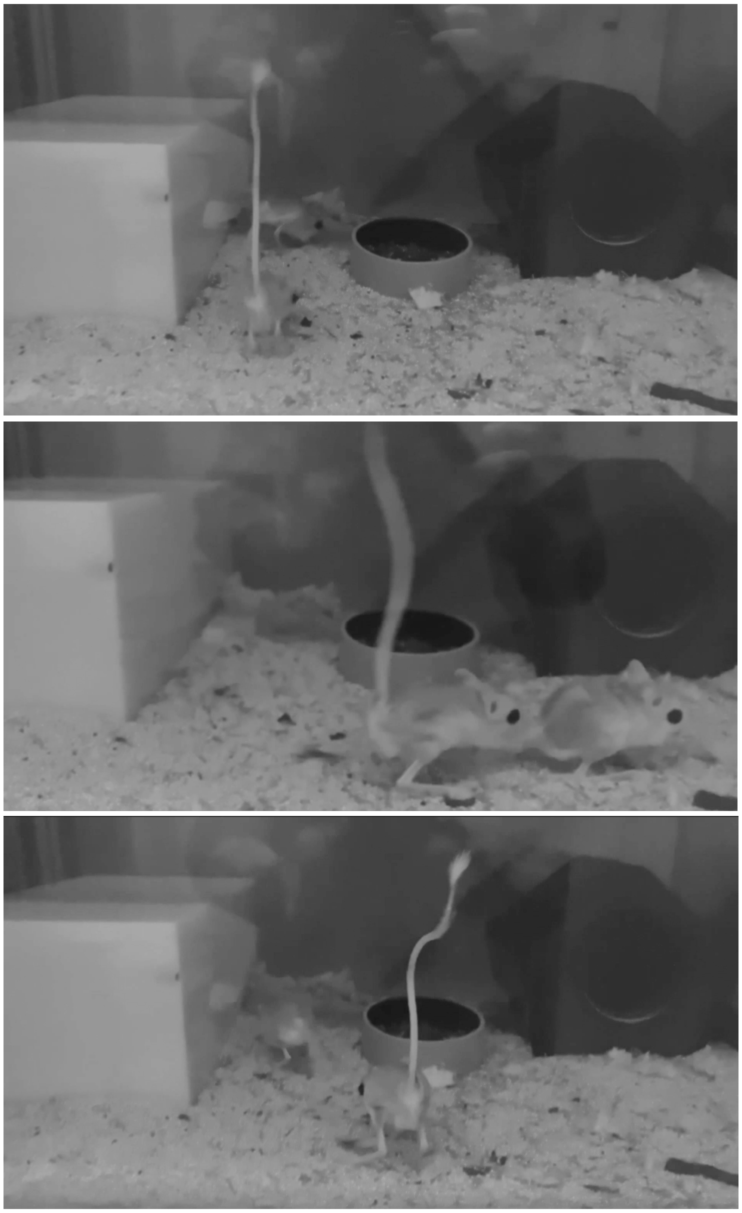
